# Detection of vasculogenic mimicry in equine ocular, oronasal, and genital squamous cell carcinoma

**DOI:** 10.1101/2025.07.05.663314

**Authors:** Sophie Schwarz, Stefan Kummer, Andrea Klang, Ingrid Walter, Barbara Nell, Sabine Brandt

## Abstract

Squamous cell carcinoma (SCC) is the most common malignant tumor disease in horses. It predominantly affects the ocular, oronasal, and anogenital region. Equine SCC is difficult to treat, also because important aspects of SCC development and metastasis are still unclear. We previously provided evidence that equine SCC cells can adopt a stem cell-like phenotype as a hallmark of malignant progression. Here, we investigated whether equine SCCs harbor endothelial-like tumor cells that form an alternative network of pseudo-vessels better known as vasculogenic mimicry (VM). Following histopathological diagnosis, 43 equine SCCs or precursor lesions (15 ocular, 14 genital, 14 oronasal tumors) were PCR-screened for equine papillomavirus (EcPV) infection. Subsequently, formalin-fixed paraffin-embedded-sections of all tumors were analyzed by Periodic Acid-Schiff (PAS) reaction and immunohistochemical (IHC) staining for endothelial cell marker CD31. Obtained micrographs were evaluated by a scientific board to unanimously identify sections of intact tumor tissue displaying PAS-positive, CD31-negative lumens harboring erythrocytes. Thirteen lesions exhibiting these features were subjected to triple immunofluorescence (IF) staining for CD31, pan-cytokeratin (KRT) and type 4 collagen (Col4) or alpha smooth muscle actin (αSMA) to confirm the presence of VM, and to determine whether pericytes have a role in this phenomenon. All genital and 50% of oronasal lesions scored positive for EcPV type 2, whilst ocular lesions tested negative. One mandibular SCC harbored EcPV type 5. Six genital, three oronasal, and four ocular tumors unambiguously exhibited VM as revealed by CD31-/PAS+ vessel-like structures containing erythrocytes, the detection of CD31-negative cells lining their lumens, and the presence of Col4 and αSMA in this lining. Detection of these two proteins in the context of VM suggests that VM-forming cancer cells recruit pericytes to enhance channel formation and stability. To our knowledge, this is the first report providing evidence of VM in equine cancer, and more generally, SCC in animals.

## Introduction

Squamous cell carcinoma (SCC) is a malignant tumor disease in vertebrates that arises from epithelial keratinocytes. In humans, SCCs can develop at any site of the skin, and likewise affect mucosal sites such as the cervix or the head-and-neck (HN) region. Interestingly, 100% of cervical carcinomas, more than half of anogenital SCCs, and up to 50% of HNSCCs are induced by high-risk human papillomaviruses (hrHPVs)[1]. HrHPV-unrelated lesions develop following excessive exposure to damaging factors such as UV-radiation, tobacco, or alcohol, and in immunosuppressed individuals [2–7].

In horses and other equid species, SCCs mainly occur in the ocular, oronasal, and anogenital region [8, 9]. Disease onset is characterized by the development of precursor lesions, i.e., actinic keratosis, whitish plaques, and/or papillomata, that progress to carcinomas in situ (CIS), and ultimately to SCCs [9]. Early diagnosis is key to effective therapy that usually consists of the excision of affected tissue with wide surgical margins. In case of advanced ocular or penile SCC, disease is usually treated by the exenteration of the affected eye, or the en-bloc resection of the penis [10–12]. Oronasal lesions often remain unnoticed until they cause obvious symptoms such as respiratory problems, ataxia, osseous deformation, and emaciation, leading to the euthanasia of the affected animal [10].

In 2010, we reported on the identification of a novel equine papillomavirus (EcPV) termed EcPV2 from genital CIS and SCCs [13]. Today, there is ample evidence that EcPV2 causes virtually 100% of equine genital SCCs [13, 14]. In addition, 20 to 50% of equine oronasal carcinomas were shown to harbor EcPV2 DNA, suggesting an etiological association of the virus with a subset of equine HNSCCs, similar to what is reported for HPV16 in relation to human HNSCC [15, 16]. In contrast, ocular SCCs are usually EcPV2-free, indicating that the disease is promoted by other factors [14, 15]. These include UV radiation [9] in combination with predisposing genetic factors such as the missense mutation in the damage-specific DNA-binding protein 2 (DDB2) identified in horses of Haflinger breed [17–19].

It is widely accepted today that the high plasticity of cancer cells represents a major hallmark of tumor growth and metastasis in human disease [20–22]. Epithelial tumor cells can acquire mesenchymal properties via a biological program termed epithelial mesenchymal transition (EMT). This transition endows tumor cells with the capacity to detach from the primary lesion, migrate through the extracellular matrix, and enter the vasculature. Once they have reached a secondary site, they extravasate, revert to epithelial-type cells by mesenchymal-endothelial transition (MET) and settle down as metastasis [23–25]. Tumor cells can also switch to a stem cell-like phenotype also known as cancer stem cells (CSCs) [22, 26–29]. Alike all stem cells, CSCs are long-lived, able to self-renew, and to differentiate into various cell types including endothelial-like cancer cells [30].

In 1999, Maniotis and colleagues were the first to describe the formation of an alternative network of micro-channels by endothelial-like melanoma cells *ex vivo* and *in vitro*. This network was termed vasculogenic mimicry (VM) as it closely resembled normal vasculature, contained erythrocytes, and was thought to assure tumor perfusion [31]. Meanwhile, VM has been identified in various human cancers including SCCs [31–36]. Importantly, VM promotes tumor cell dissemination, thus being associated with poor prognosis [36]. In addition, VM networks are resistant to anti-angiogenic therapy as they do not depend on classical pro-angiogenic growth factors and cytokines required for the formation of normal vasculature [34, 37].

In animals, current information on VM is only scarce, and limited to mammary cancer, melanoma, and osteosarcoma in dogs [33, 38, 39]. Given the crucial role of VM in tumor metastasis and therapy-resistance on one hand, and the total lack of knowledge regarding this phenomenon in equine cancers on the other, we analyzed a series of equine EcPV-positive and -negative SCCs/SCC precursor lesions for the presence of VM using PAS reaction and immunohistochemical staining for CD31. Subsequently, we validated our findings by triple immunofluorescence (IF) labelling for CD31, pan-cytokeratin, and type IV collagen, or alpha smooth muscle actin.

## Materials and Methods

### Sample material, tumor diagnosis and screening for EcPV

A total of 43 tumor samples were included in the study. Lesions originated from the ocular (n=15), the oronasal (n=14), and the genital region (n=14), and were collected at the University of Veterinary Medicine, Vienna, during therapeutic excision or requested necropsy with the horse owners’ written consent. Patient, disease, and sample specifications are provided in S1 Table.

The tumor samples were available as DNA aliquots and FFPE-embedded material. The histopathological diagnosis of SCC (n=40) or carcinoma in situ (CIS; n=3) was made from hematoxylin-eosin (HE) stained FFPE sections as described previously [40]. Tumor DNA was quality-checked and tested for the presence of ECPV type 2, 3, and 5 DNA as published previously [40].

### Immunohistochemical staining for CD31 and Periodic Acid Schiff reaction

Sections (2.5 µm) from the 43 tumors included in the study, and from equine kidney (positive control) were stained for the endothelial cell marker CD31 and by Periodic Acid Schiff (PAS) reaction to discern normal microvessels surrounded by CD31-positive endothelial cells from VM, i.e., channel-like structures predominantly lined by CD31-negative cancer cells. Intraluminal erythrocytes were detected on basis of their morphology, localization, and slight PAS staining. Normal blood vessels in the lesions served as internal positive control; negative control sections were obtained by omission of the primary antibody (no-primary-antibody controls; npac).

In a first step, sections were deparaffinized with xylene, and successively dehydrated in 100%, 96%, and 70% ethanol. Then, sections were treated with 0.3% H_2_O_2_/methanol to block peroxidase activity. Heat-induced epitope retrieval (HIER) was performed in 0.1 M citrate buffer (pH 6) for 30 min in a steamer at 94–100 °C. Following rinsing in phosphate buffered saline (PBS), sections were blocked with 1.5% normal goat serum (Dako, Glostrup, Denmark) to minimize unspecific antibody binding.

In a next step, sections were incubated with rabbit anti-human CD31 antibody (Ab; 1:1000 in PBS; Cell Marque, Sigma-Aldrich, Vienna, Austria) at 4°C overnight. Following washing with PBS, sections were incubated with poly-horseradish peroxidase (HRP)-conjugated anti-rabbit Ab (ready to use; BrightVision, ImmunoLogic, Duiven, The Netherlands) for 60 min at room temperature (RT). Then, Ab-bound protein was incubated with diaminobenzidine (DAB) chromogen (ThermoFisher Scientific, Vienna, Austria) for 5 min at room temperature. After rinsing in sterile water, sections were subjected to PAS reaction by oxidization in Periodic acid solution (0.5%) for 20 min, triple rinsing in distilled water, incubation in Schiff’s reagent (Morphisto GmbH, Offenbach am Main, Germany) and rinsing with water. Then, sections were counter-stained in Mayer’s hematoxylin (Morphisto GmbH), rinsed with water, and dehydrated as described above. Finally, slides were mounted with Epredia™ Consul-Mount™ (ThermoFisher Scientific).

For detection of VM, sections were evaluated for the presence of PAS-positive, CD31-negative lumina exhibiting a meandering pattern using a Zeiss Axio Imager Z2 microscope (Zeiss, Oberkochen, Germany). Images were taken with an AxioCam MRc5 camera (Zeiss) at 40x magnification and ultimately presented to the board of authors to identify tumors that unambiguously displayed VM.

### Triple immunofluorescence (IF) staining for CD31, pan-cytokeratin, and type IV collagen

IF staining was carried out from sections of 13 selected tumors convincingly exhibiting VM to further confirm this finding. Sections (2.5-µm) of the lesions, and equine esophagus (positive control) were triple-stained for CD31, pan-cytokeratin (KRT), and type IV collagen (Col4). To this end, the sections were deparaffinized, rehydrated, subjected to HIER, and blocked exactly as described above. In a next step, sections were incubated with the first primary Ab, i.e., mouse anti-human pan-KRT Ab (1:500 in PBS; Abcam Ltd., Cambridge, UK) at 4°C overnight. After washing and incubation with polyHRP-conjugated goat anti-mouse Ab (ready to use; BrightVision, Immunologic) for 60 min at RT, the signal was developed with an Alexa 568-labeled tyramide solution (ready to use; Invitrogen, ThermoFisher Scientific; orange signal) for 10 min at RT. After elution of the Abs with 0.01 M citrate buffer (pH6) for 60 min at 95°C in a water bath, and blocking of unspecific Fc-receptors, the sections were incubated with the second primary Ab, i.e., rabbit anti-human CD31 Ab (1:500 in PBS; Cell Marque) at 4°C overnight. After washing and incubation with polyHRP-conjugated goat anti-rabbit Ab (ready to use; BrightVision, Immunologic) for 60 min at RT, the signal was developed with an Alexa 647-labeled tyramide solution (ready to use; Invitrogen, ThermoFisher Scientific; red signal) for 10 min at RT. Following Ab elution and blocking as described above, the sections were incubated with the third primary Ab, i.e., mouse anti-human Col4 Ab (1:100 in PBS; Cell Marque) at 4° overnight. Then, binding of this third primary Ab was detected by incubation with polyHRP-conjugated goat anti-mouse Ab (ready to use; BrightVision, Immunologic) for 60 min at RT and subsequent development with Alexa 488-labeled tyramide solution (ready to use; Invitrogen, ThermoFisher Scientific; green signal) for 10 min at RT. Finally, cell nuclei were counterstained with DAPI (Sigma Aldrich, St. Louis, MO, USA), and the sections were mounted with Aqua-PolyMount (Polysciences, Szabo-Scandic, Vienna, Austria). Stained sections were digitized using a slide scanner (VS200, Evident Europe GmbH, Hamburg, Germany) equipped with a 20x lens. Npac sections were included in all reactions.

### Triple IF staining for CD31, pan-cytokeratin, and alpha smooth muscle actin

Finally, staining for alpha smooth muscle actin (α-SMA; ACTA2) in combination with CD31 and pan-KRT was carried out. Sections of the 13 selected tumors with VM, and of equine esophagus as positive control were analysed using the triple-staining protocol described above with the only difference that sections were stained in a different order: First for α-SMA using a mouse anti-human SMA Ab (1:1000 in PBS; Dako), the polyHRP-conjugated goat anti-mouse Ab (ready to use; BrightVision, Immunologic) as secondary Ab, and Alexa 488-labeled tyramide solution (Invitrogen, ThermoFisher Scientific; green signal); then for pan-KRT (with Alexa 568; orange signal), and lastly for CD31 (with Alexa 647; red signal) exactly as described above. In a final step, sections were counterstained with DAPI, mounted, and digitized as specified above.

## Results

### Distribution of EcPV infection among lesions

DNA from the 40 equine SCCs and three CIS was screened for infection by carcinogenic EcPV types 2, 3, and 5 [40]. From the 14 oral SCCs, seven scored positive for EcPV2, and one for EcPV5 DNA. In contrast, only one periocular SCC tested positive for EcPV2, whilst the 14 ocular lesions tested negative. All genital tumors scored positive for EcPV2 DNA (S1 Table). None of the lesions contained EcPV3 DNA. The negative, positive, and no-template controls included in all reactions yielded the expected results.

### CD31/PAS-staining reveals VM-like structures in a subset of lesions

Following EcPV PCR of tumor DNA isolates, FFPE-sections from the 43 lesions were subjected to CD31-staining and PAS reaction. Images were subsequently evaluated by the scientific board of authors for the the presence of VM. Sections of 13 lesions (30.23%) unambigously exhibted hollow structures that were mainly or exclusively lined by CD31-negative cells and contained erythrocytes (Table 1; Fig 1). Sections of the other 30 lesions (60.77%) tested negative for VM.

**Fig 1.**
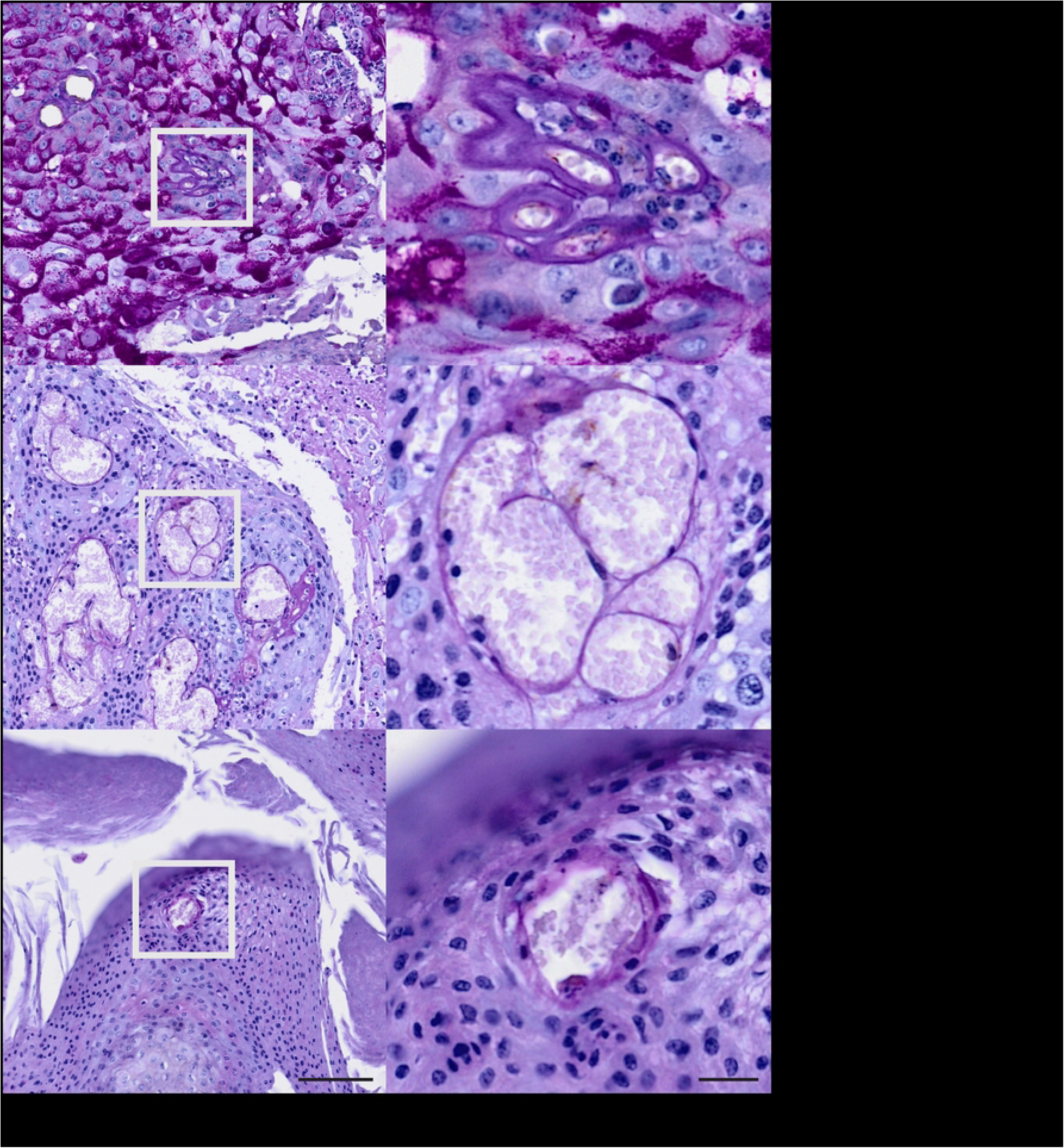
Evidence of VM in equine ocular, oropharyngeal and penile SCC. CD31/PAS-staining of tumor sections provided evidence of VM in a minimum of 13 lesions as exemplarily shown for an ocular (BOD), an oropharyngeal (JON) and a penile SCC (ERI). Staining revealed hollow structures lined by CD31-negative tumor cells (no brown signal; for controls see S1 Fig) and harboring erythrocytes as revealed by their morphological appearance and slight PAS staining). White frames depict the areas also presented at higher magnification.

**Table 1:** Horses with ocular, oronasal, and genital tumors displaying VM.

| ID | Breed | Age | Color | Sex | Diagnosis | EcPV type |
| --- | --- | --- | --- | --- | --- | --- |
| BEL | Haflinger | 14 | sorrel | M | SCC of the cornea OD (resected) | N |
| BOD | WB | 19 | chestnut | G | invasive SCC, nictitating membrane OD (resected) | N |
| MON | Connemara | 15 | grey | M | 2012: SCC OS, exenteration; 2014: metastases (†) | 2 |
| DEX | Trakehner | 16 | piebald | G | CIS nictitating membrane OD (resected) | N |
| JON | Pony | 25 | chestnut | G | SCC gingiva, palate, pharynx (†) | 2 |
| MIS | Haflinger mix | 13 | bay | G | SCC nasal cavity, concho-frontal sinus (excised) | N |
| SNI | Connemara | 16 | grey | M | SCC mandible, LN metastases (†) | 5 |
| ARO | Hanoverian | 18 | bay | G | invasive SCC penis, inguinal LN metastases (†) | 2 |
| BLÁ | Icelandic | 23 | chestnut | G | SCC penis (penile en-bloc resection) | 2 |
| BOG | WB | 28 | black | G | SCC penis, inguinal LN metastasis (†) | 2 |
| ERI | WB | 15 | bay | G | SCC glans penis (partial penile resection) | 2 |
| GJE | WB | 20 | skewbald | M | SCC vulva (resection) | 2 |
| MLK | Cob | 15 | piebald | G | large SCC penis (penile en-bloc resection) | 2 |

ID: horse identification code; Age: in years; G: gelding (castrated male); M: mare (female); SCC: squamous cell carcinoma; CIS: carcinoma in situ; LN: lymph node; OD: oculus dexter, right eye; OS: oculus sinister, left eye; in parentheses: outcome, with † designating euthanization; N: no EcPV DNA detected.

### Triple IF staining for CD31, KRT, and Col4 confirms the presence of VM in equine SCCs/CIS

In a next step, sections of the 13 tumors with evidence of VM according to CD31/PAS-staining were subjected to triple IF labeling for CD31 (red signal), pan-cytokeratin (KRT; orange signal), and Col4 (green signal). Stained normal vasculature versus VM in sections of an ocular (BOD; Fig 2A), an oropharyngeal (JON; Fig 2B), and a penile SCC (ERI, Fig 2C) are exemplarily shown in Fig 2. Additional data are provided in S2 Figs A-C. As expected, normal blood vessels in the tumors were clearly lined by CD31-positive endothelial cells and surrounded by a Col4-positive membrane. Tumor cells stained KRT-positive in agreement with their epithelial origin (Fig 2A-C, top rows). Alike normal vasculature, VM channels contained erythrocytes as detected by their autofluorescence, and exhibited a Col4-positive lining. Yet, they clearly scored CD31-negative (Fig 2A-C, bottom rows), as best illustrated by the black-and-white image representing the red channel (Fig 2A-C, bottom row, CD31).

**Figs 2.**
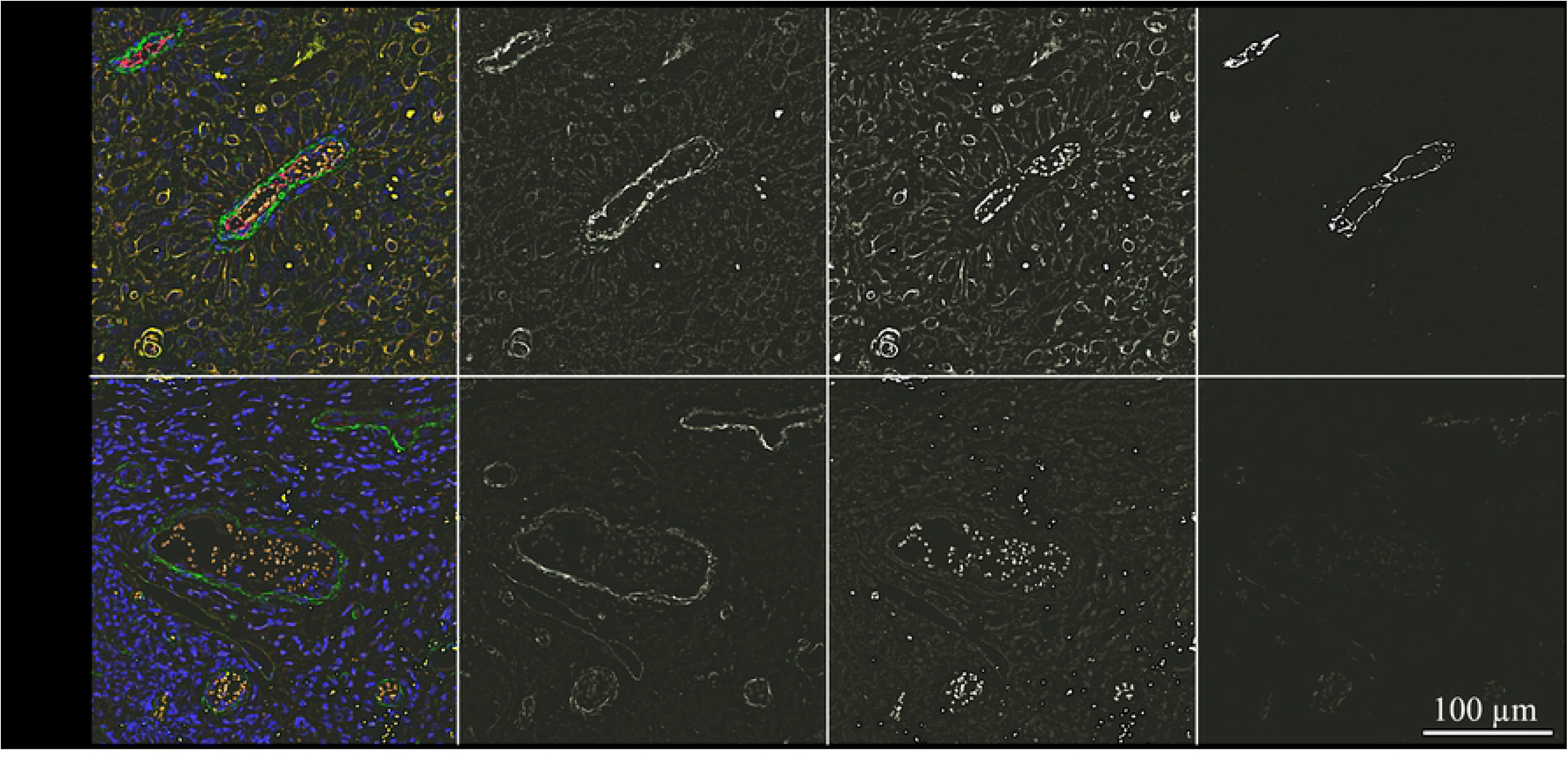
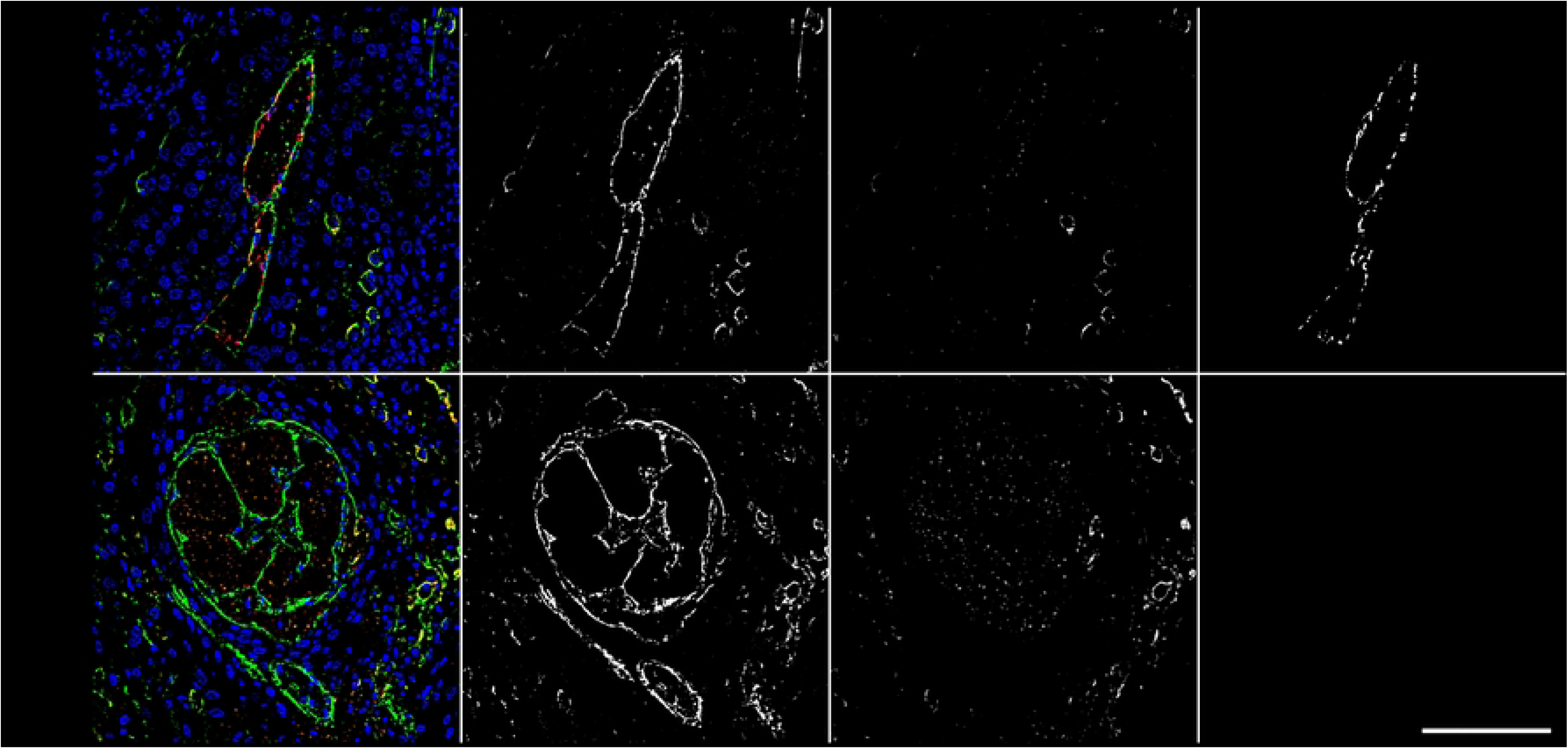
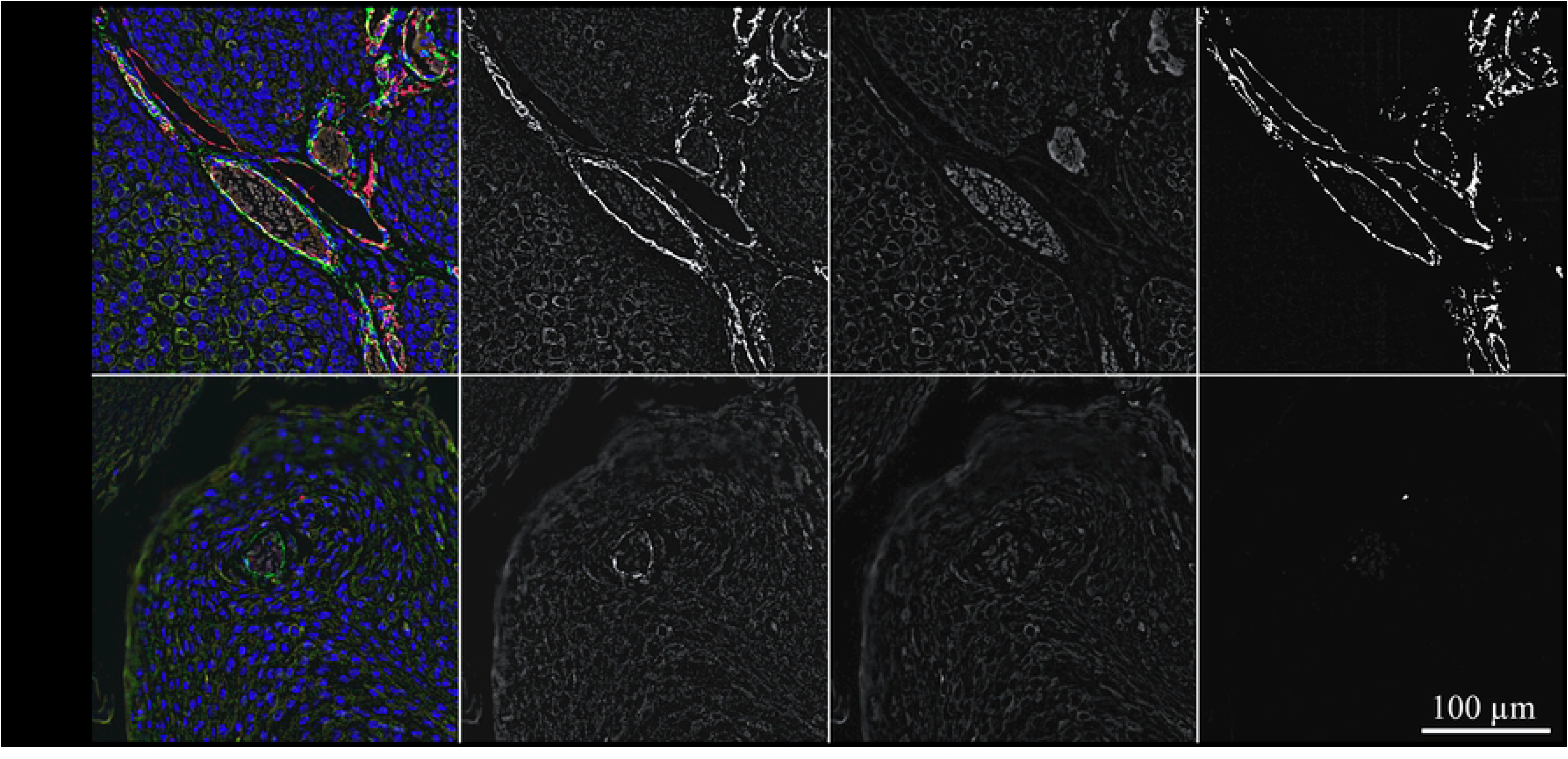
A-C. Intralesional presence of VM shown by CD31/KRT/Col4 IF labeling. Figures 2A-C depict sections of an ocular (2A; BOD), an oropharyngeal (2B; JON), and a penile SCC (2C; ERI) IF-stained for CD31 (red signal), KRT (orange signal), and Col4 (green signal). The blue signal reflects DAPI-stained cell nuclei. Individual staining results are presented in black and white in addition to merged images from normal blood vessels and VM structures to allow for direct comparison and optimum signal evaluation. In some cases, VM structures displayed a mosaic pattern, with a few CD31-positive cells in the lumens’ lining. Fig 3 exemplarily illustrates this observation.

**Fig 3.**
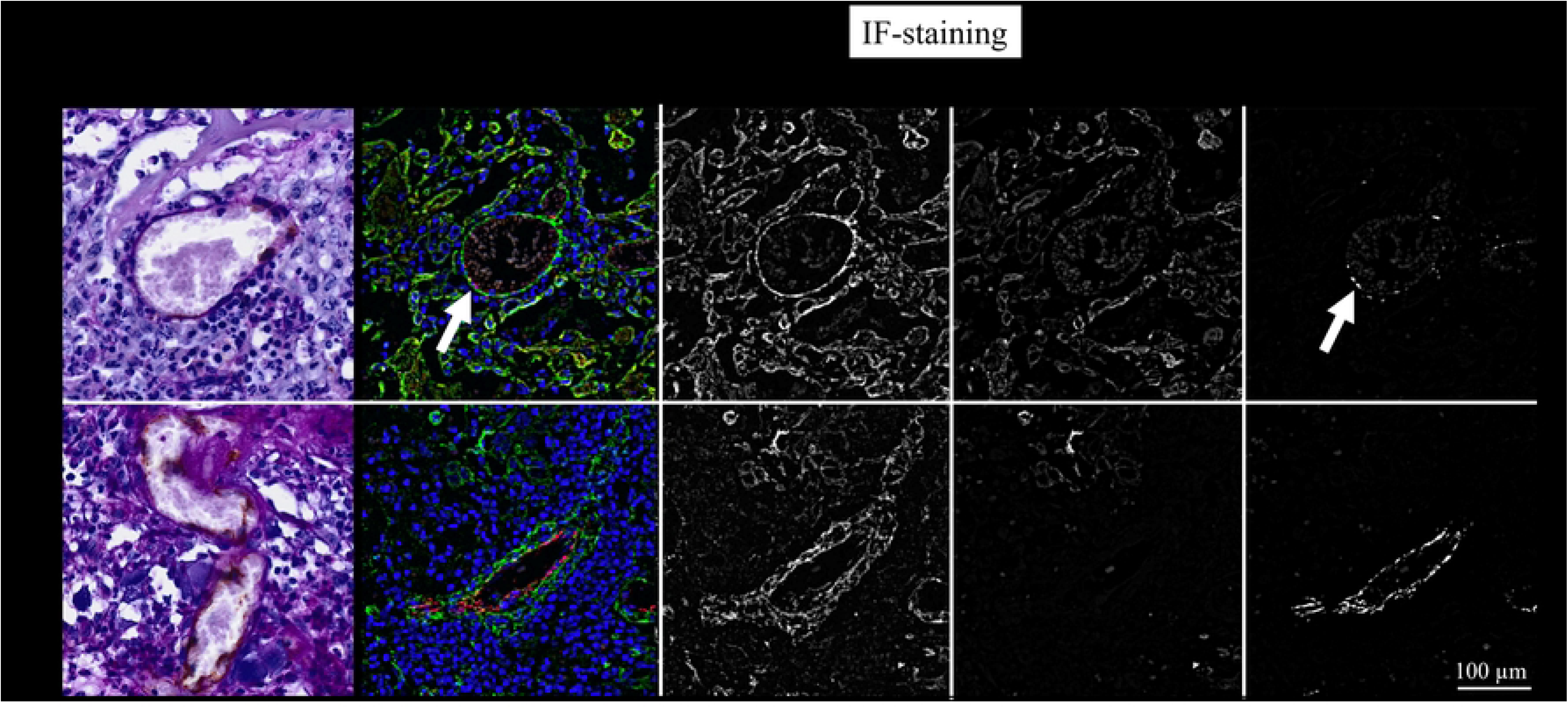
Mosaic pattern of VM observed in sections of an ocular SCC. The figure depicts CD31/PAS and CD31/KRT/Col4 IF stained sections of an ocular SCC (MON) exhibiting VM (top row) or normal blood vessels (bottom row). CD31/PAS staining allowed for distinction of pseudo-vessels delimited by tumor cells from normal blood vessels exhibiting a CD31+ endothelial cell lining. Of note, the VM lining also harbored a few CD31+ cells. This finding was confirmed by triple IF staining (bottom row) for CD31 (red signal), KRT (orange signal), and Col4 (green signal). The figure shows merged IF images from normal blood vessels and VM structures for direct comparison, with white arrows pointing to CD31+ cells in VM. Individual staining results presented in black-and-white allowed for optimum signal evaluation.

### Triple IF staining for CD31, KRT, and α-SMA suggests that pericytes are involved in VM

In a final step, sections of the 13 tumors with evidence of VM were IF-stained for CD31 (red signal), pan-cytokeratin (KRT; orange signal), and α-SMA (green signal). Labeling results are exemplarily presented in Figs 4A-C showing stained sections of an ocular (Fig 4A), an oronasal (Fig 4B) and a penile SCC (Fig 4C). Additional data are provided in S2 Figs A-C.

**Figs 4.**
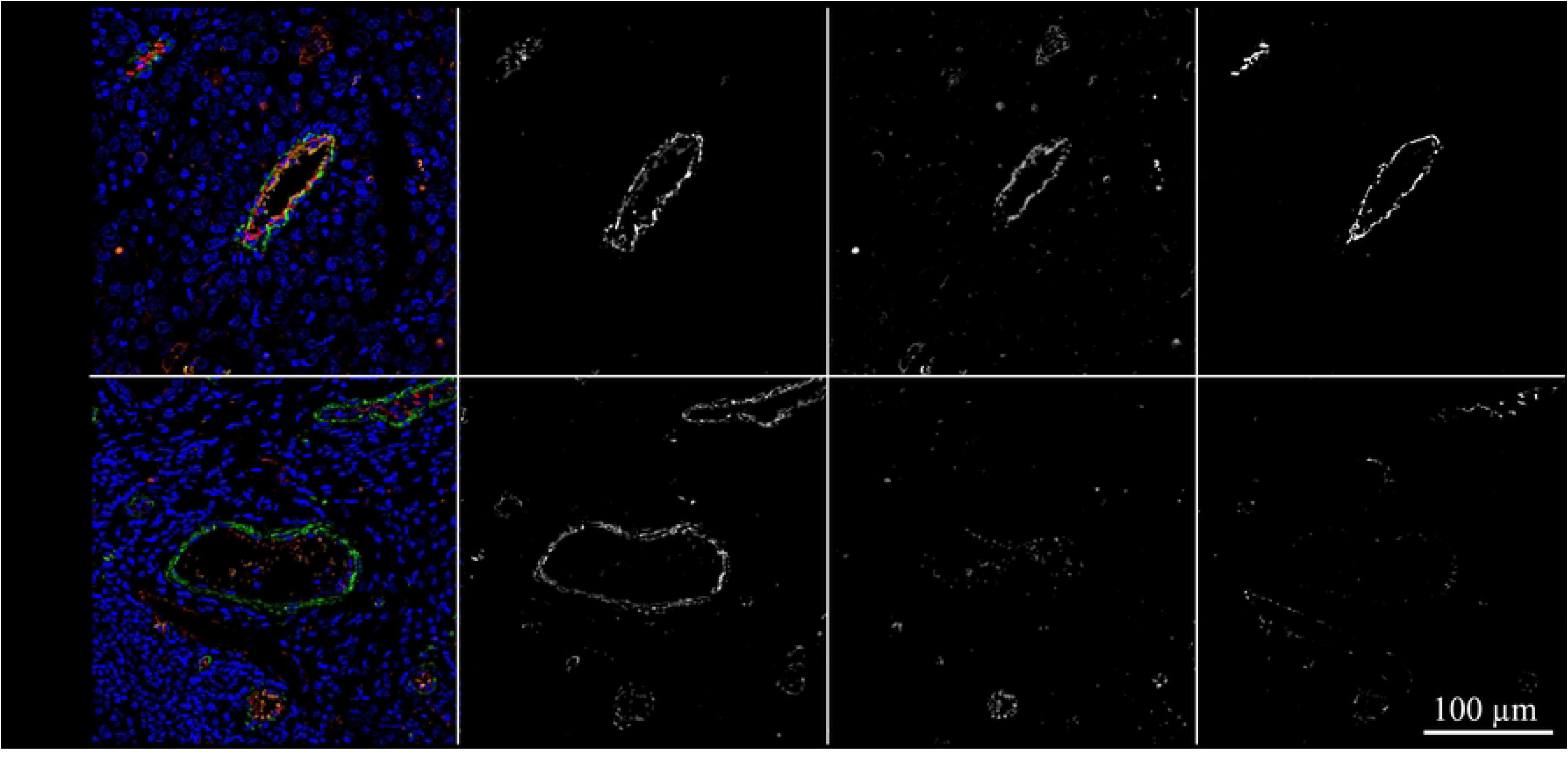
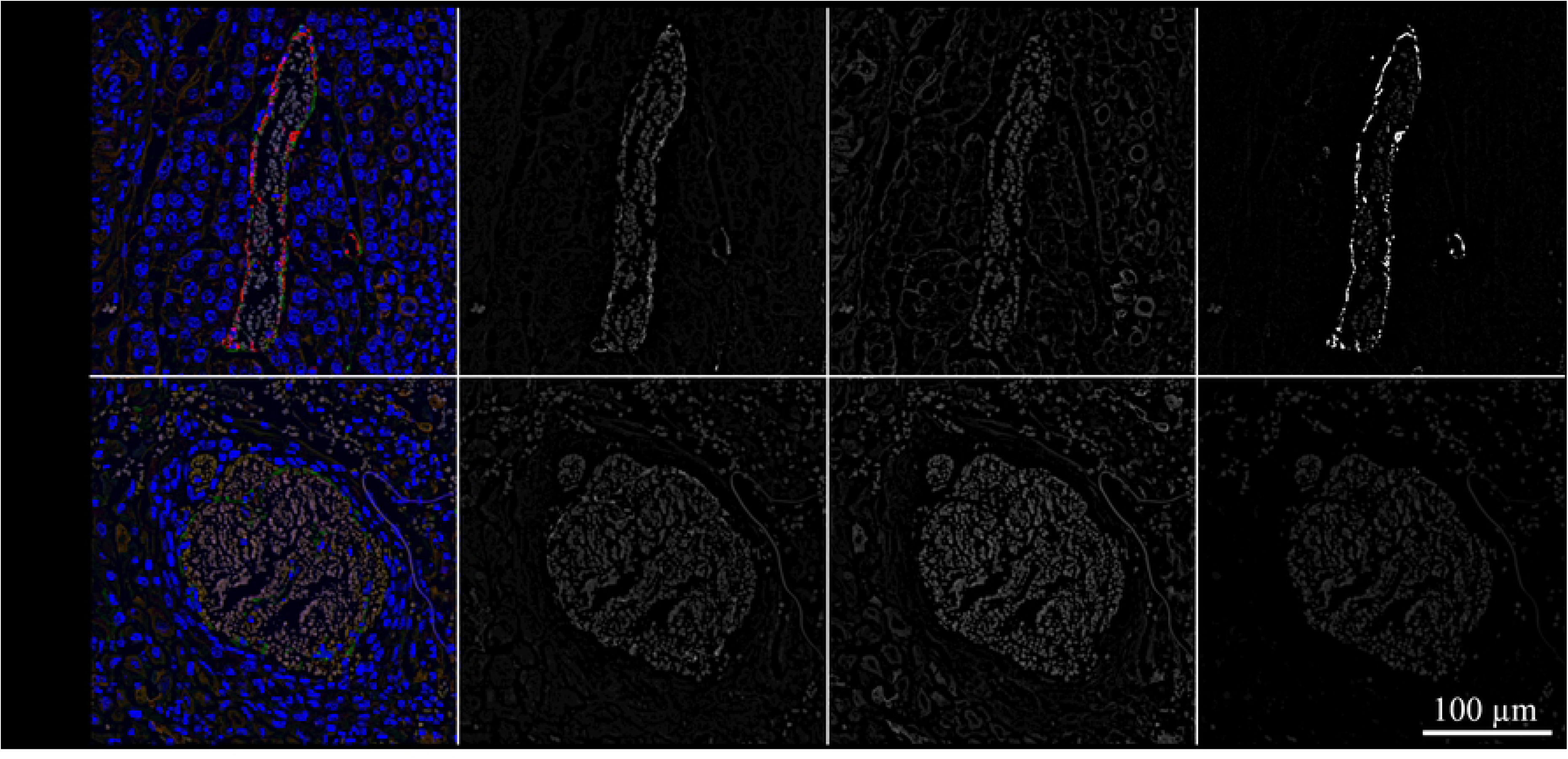
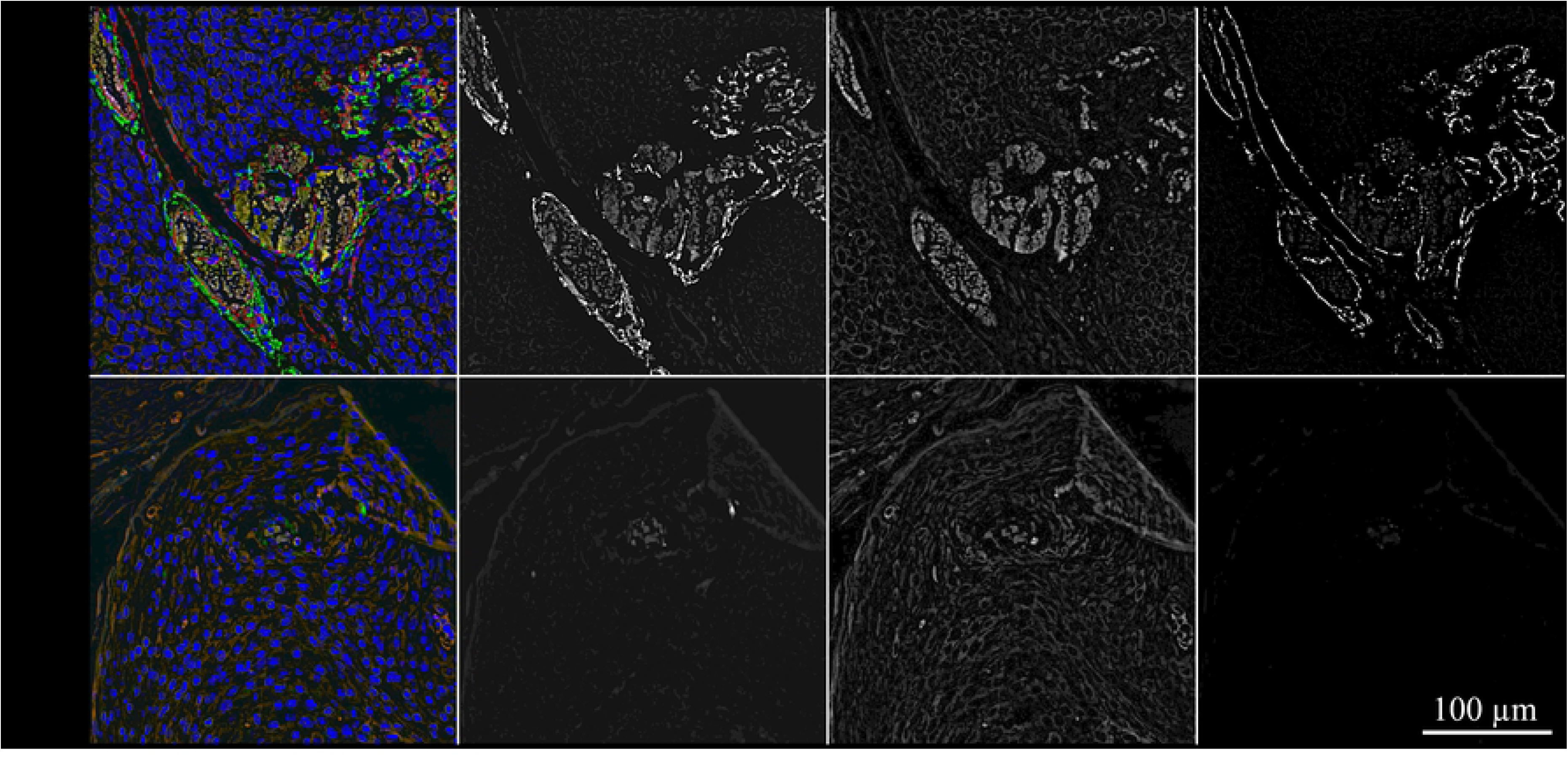
A-C: The lining of VM channels also harbors pericytes. Figs 4A-C depict sections of an ocular (4A; BOD), an oropharyngeal (4B; JON), and a penile SCC (4C; ERI) that were IF-stained for CD31 (red signal), KRT (orange signal), and α-SMA (green signal). The blue signal reflects DAPI-stained cell nuclei. The figures show merged images from normal blood vessels and VM structures for direct comparison. In addition, individual staining results are presented in black-and-white to allow for optimum signal evaluation.

The lining of normal blood vessels and VM channels both displayed α-SMA labelling, albeit with varying intensity as shown in Figs 4A-C and S3 Fig. Normal blood vessels showed more intense α-SMA labelling in comparison with suspected VM throughout the tumor sections. It has to be noted that the green signal exhibited by α-SMA positive cells was partly overlayed by the red CD31 signal emitted by endothelial cells as illustrated in Fig 4B.

## 4. Discussion

The ability of tumor cells to adopt an endothelial cell-like phenotype that builds an alternative network of pseudo-vessels, i.e., VM, is tightly linked to the metastatic behavior of cancers including SCCs [24, 41, 42]. In human HNSCC, VM is relatively well documented and the mechanisms underlying VM formation are beginning to be elucidated [36,44–48].

In equine oncology, the field of VM is completely unexplored. Armando et al., and our group have previously shown that equine EcPV2-positive and -negative SCC cells can undergo (partial) EMT and also switch to a CSC phenotype [40, 43], as similarly reported for human HPV^+/-^ SCC cells [22, 28, 29]. In the herein reported work, we assessed equine SCCs or CIS for EcPV infection, and subsequently for the presence of VM structures built by tumor cells.

As expected on the basis of previous findings [14, 16], 100% of genital SCCs or CIS, and 50% of oronasal SCCs tested positive for EcPV2 DNA, whilst virtually all ocular lesions scored EcPV-negative. This finding corroborates the assumption that EcPV2 is actively involved in the development of anogenital SCC, and a subset of oronasal SCCs [14, 16]. Simultaneously, it strengthens the concept that ocular SCC constitutes a different tumor entity with respect to its etiology [9, 19, 44, 45].

Following EcPV screening, tumor sections were subjected to a series of immunostaining reactions using primary mouse or rabbit anti-human antibodies. To rule out the possibility of false positive or false negative results, different antibody clones per target were carefully evaluated in a pilot study using control tissue according to the Human Protein Atlas (https://www.proteinatlas.org). This led to the identification of anti-human antibodies with maximum target specificity also in the horse. To reach maximum technical accuracy, a positive control (equine kidney or esophagus) and no-primary-antibody controls were included in all staining experiments and yielded expected results.

The presence of VM was first addressed using PAS reaction in combination with IHC staining for the endothelial cell marker CD31. This method is deemed suitable for the detection of VM as it allows the visualization of erythrocyte-containing lumens displaying a PAS-positive, but CD31-negative lining [31, 46, 47]. Obtained micrographs were subsequently evaluated in a board meeting of all authors. VM was identified according to the above-mentioned standard criteria. In addition, the board aimed at ascertaining that detected VM regions truly represented functional channels by excluding tumors exhibiting signs of denaturation, necrosis, or glycoprotein aggregates. This led to the selection of thirteen lesions exhibiting VM according to PAS/CD31 staining.

In a next step, sections of these selected tumors were subjected to triple IF labeling for CD31, the squamous epithelial cell marker pan-cytokeratin (KRT), and type IV collagen (Col4) as a substitute for PAS. Micrographs were taken from intralesional blood vessels and VM regions to allow for comparative evaluation. Intraluminal erythrocytes were detected by their autofluorescence that was most pronounced in the red color spectrum. Normal blood vessels differed from VM in that they were clearly lined by CD31-expressing endothelial cells and that the lining also exhibited a more pronounced Col4 signal. VM structures were surrounded by cells that usually scored negative for CD31, thus exhibiting a tubular VM pattern (48). Interestingly, some VM linings also contained a limited number of CD31-positive cells, thus possibly exhibiting a mosaic pattern. This pattern could not be unambiguously identified by PAS/CD31-staining, emphasizing that a combination of this method with IHC- or IF-staining for additional markers is better suited for the identification of VM, as already suggested previously [48–50]. The exact nature of these CD31-expressing cells in the VM-lining is unclear. When mosaic VM patterns were first observed, the CD31-positive cells in the lining were interpreted as normal endothelial cells located at the junctions between VM channels and normal blood vessels [51]. Meanwhile, there is a growing body of evidence that CSCs can differentiate into endothelial cancer cells that express endothelial cell-specific proteins including CD31 [32, 48, 52–55]. If this is the case, the CD31-positive cells occasionally detected in the VM lining of equine SCCs would correspond to tumor cells. On the other hand, the immunoglobulin-like receptor CD31 is not only expressed by endothelial cells, but also by immune cells, including T- and B-lymphocytes as well as dendritic cells (DCs)[56]. On these grounds, we cannot rule out the possibility that CD31^+^ cells occasionally detected in the lining of VM may correspond to leukocytes. Hence, more work is needed to conclusively explain the CD31 expression pattern observed in a subset of VM.

Equine SCCs were previously shown to harbor tumor cells corresponding to a hybrid epithelial-mesenchymal phenotype [40]. This finding was not surprising, since complete EMT is only achieved *in vitro*. *In vivo*, human SCC cells usually undergo partial EMT (pEMT), i.e., they acquire mesenchymal properties allowing for example migration, but also preserve epithelial features. This hybrid status is thought to confer an advantage to the cancer cells [57, 58]. Similarly, CSCs may not fully differentiate to endothelial cancer cells, but keep CSC features. This assumption is strengthened by the finding that endothelial-like cancer cells can also express stem cell-specific proteins [48]. The degree to which CSCs acquire endothelial cell-specific features may eventually depend on (micro-)environmental factors that remain to be fully elucidated.

The presumed transition of CSCs into “endotheloid” cancer cells that form VM structures also provides an explanation for the poor KRT expression observed in the VM linings in equine SCCs. In humans, the intermediate filament of endothelial cells is composed of vimentin, reflecting their mesenchymal origin. KRT expression by endothelial cells is rather confined to lower animal species e.g., of the classes Amphibia or Pisces [59, 60]. On these grounds, it is conceivable that the equine endotheloid tumor cells composing the immediate VM lining do not express high levels of cytokeratins. In a final experiment, the thirteen selected tumors were IF-labeled for α-SMA in combination with CD31 and KRT. The rationale for this approach was that α-SMA is expressed by smooth muscles, but also pericytes, which are suspected to have a major role in VM. In line with the above-mentioned concept that CSCs can differentiate into endothelial cancer cells, Thijssen and colleagues provided evidence that VM-associated (“VM^+^”) cancer cells (i.e., endothelial cancer cells) can “phenocopy” important steps of normal angiogenesis in that they can recruit pericytes for the construction and stabilization of the VM network [34].

Triple IF labeling of tumor sections and VM regions for CD31, KRT, and α-SMA resulted in all intralesional blood vessels exhibiting an α-SMA-positive lining. This finding was anticipated on the grounds that this protein is ubiquitously expressed in normal vessel networks [61]. Importantly, the majority of VM regions analyzed also displayed this feature, although not being surrounded by smooth musculature [31]. This interesting observation supports the idea that endotheloid cancer cells hijack pericytes for the creation of an independent network of stable pseudo-vessels assuring the supply of the tumor with oxygen and nutrients, and the dissemination of tumor cells [34].

Under physiological conditions, the recruitment of pericytes by endothelial cells also results in the maturation of blood vessels, as it induces the expression of core matrix proteins such as laminin, and type IV and VI collagens that form the matrix of the endothelial basement membrane [62]. Detected Col4 expression not only in the lining of intralesional blood vessels, but also in VM walls is a further indication of pericyte recruitment by endotheloid equine cancer cells.

## Conclusions

This study is the first to provide evidence of VM in equine SCC and CIS. Intralesional structures exhibiting the characteristic features of VM, i.e., lumens containing erythrocytes and lined by predominantly CD31-negative cells, were detected irrespective of the EcPV infection status and the tumor site (ocular, oronasal, or genital).

All tumors convincingly displaying VM were late-stage or recurrent lesions, with metastases being commonly observed. This finding agrees with the concept that VM is associated with advanced, metastasizing disease. However, mild-type SCC precursor lesions such as intraepithelial neoplasia, or papilloma, were not included in the study. Hence, the onset of VM at earlier stages of disease cannot be ruled out. Interestingly, our data also indicate that the tumor cells recruit pericytes to maintain nutrient supply by stabilizing vascular-like structures in tumor tissue. Further work is needed to fully elucidate the role of pericyte recruitment in VM in equine SCCs.

## Supporting information: Figure legends

**S1 Fig. CD31- and PAS-staining of an equine kidney section (positive control) highlights the specificity of the reactions.**

The figure depicts CD31-positive capillaries (brown signal) in equine kidney tissue including a glomerulus. Pas-staining (magenta) highlights the basement membranes of the glomerular capillary loops and the tubular epithelium.

**S2 Fig A-C. VM in equine SCC as revealed by triple IF labeling for CD31, KRT, and Col4 or ⍺-SMA.**

The figures depict sections from ocular (A), oronasal (B), and penile SCCs (C) that were IF-stained for CD31 (red signal), KRT (orange signal), and Col4 or α-SMA (green signal). The blue signal reflects DAPI-stained cell nuclei. The figures show merged images from normal blood vessels and VM structures for direct comparison.

**S3 Fig. Triple IF-staining of equine esophagus sections as control underscores the specificity of the reactions.**

Equine esophagus sections were IF-stained for CD31 (red signal), KRT (orange signal), and Col4 (left side; green signal), or α-SMA (right side; green signal). The figures show merged images and the no-primary-antibody controls in color, and individual staining results in black-and-white. The blue signal reflects DAPI-stained cell nuclei. Left-side-images display the esophageal mucosa lined by squamous epithelium (KRT^+^; orange signal), the submucosa, and a part of the muscular layer. Ride-side-images show blood vessels in the submucosa at higher magnification. They are lined by CD31^+^ endothelial cells (red signal) surrounded by the α-SMA^+^ and Col4^+^ tunicae (green signals). Scale bars = 100 µl.

## References

1. Dayyani F, Etzel CJ, Liu M, Ho CH, Lippman SM, Tsao AS. Meta-analysis of the impact of human papillomavirus (HPV) on cancer risk and overall survival in head and neck squamous cell carcinomas (HNSCC). Head Neck Oncol. 2010;2:15.

2. Claveau J, Archambault J, Ernst DS, Giacomantonio C, Limacher JJ, Murray C, et al. Multidisciplinary management of locally advanced and metastatic cutaneous squamous cell carcinoma. Curr Oncol. 2020;27(4):e399–e407.

3. Fania L, Didona D, Di Pietro FR, Verkhovskaia S, Morese R, Paolino G, et al. Cutaneous Squamous Cell Carcinoma: From Pathophysiology to Novel Therapeutic Approaches. Biomedicines. 2021;9(2).

4. Guech-Ongey M, Engels EA, Goedert JJ, Biggar RJ, Mbulaiteye SM. Elevated risk for squamous cell carcinoma of the conjunctiva among adults with AIDS in the United States. Int J Cancer. 2008;122(11):2590–3.

5. Gutzmer R, Wiegand S, Kolbl O, Wermker K, Heppt M, Berking C. Actinic Keratosis and Cutaneous Squamous Cell Carcinoma. Dtsch Arztebl Int. 2019;116(37):616–26.

6. Kallini JR, Hamed N, Khachemoune A. Squamous cell carcinoma of the skin: epidemiology, classification, management, and novel trends. Int J Dermatol. 2015;54(2):130–40.

7. Marur S, Forastiere AA. Head and Neck Squamous Cell Carcinoma: Update on Epidemiology, Diagnosis, and Treatment. Mayo Clin Proc. 2016;91(3):386–96.

8. Strafuss AC. Squamous cell carcinoma in horses. J Am Vet Med Assoc. 1976;168(1):61–2.

9. Scott DW, Miller, W. H. Jr. . Squamous cell carcinoma. In: Scott DW, Miller, W. H. Jr. , editor. Equine Dermatology. 1 ed. St. Louis, Missouri, USA: Saunders Elsevier; 2003. p. 707–12.

10. Knottenbelt DC. Squamous cell carcinoma. In: Knottenbelt DC, editor. Pascoe’s Principles and Practice of Equine Dermatology. Second Edition ed. London, UK: Saunders Elsevier; 2009. p. 427–33.

11. Pascoe RR, Knottenbelt DC. Squamous cell carcinoma. In: Saunders WB, editor. Manual of Equine Dermatology. London, UK: Harcourt Publishers; 1999. p. 261–6.

12. van den Top JG, de Heer N, Klein WR, Ensink JM. Penile and preputial squamous cell carcinoma in the horse: a retrospective study of treatment of 77 affected horses. Equine Vet J. 2008;40(6):533–7.

13. Scase T, Brandt S, Kainzbauer C, Sykora S, Bijmholt S, Hughes K, et al. Equus caballus papillomavirus-2 (EcPV-2): an infectious cause for equine genital cancer? Equine Vet J. 2010;42(8):738–45.

14. Sykora S, Brandt S. Papillomavirus infection and squamous cell carcinoma in horses. Vet J. 2017;223:48–54.

15. Knight CG, Dunowska M, Munday JS, Peters-Kennedy J, Rosa BV. Comparison of the levels of Equus caballus papillomavirus type 2 (EcPV-2) DNA in equine squamous cell carcinomas and non-cancerous tissues using quantitative PCR. Vet Microbiol. 2013;166(1-2):257–62.

16. Sykora S, Jindra C, Hofer M, Steinborn R, Brandt S. Equine papillomavirus type 2: An equine equivalent to human papillomavirus 16? Vet J. 2017;225:3–8.

17. Bellone RR, Liu J, Petersen JL, Mack M, Singer-Berk M, Drogemuller C, et al. A missense mutation in damage-specific DNA binding protein 2 is a genetic risk factor for limbal squamous cell carcinoma in horses. Int J Cancer. 2017;141(2):342–53.

18. Lassaline M, Cranford TL, Latimer CA, Bellone RR. Limbal squamous cell carcinoma in Haflinger horses. Vet Ophthalmol. 2015;18(5):404–8.

19. Singer-Berk M, Knickelbein KE, Vig S, Liu J, Bentley E, Nunnery C, et al. Genetic risk for squamous cell carcinoma of the nictitating membrane parallels that of the limbus in Haflinger horses. Anim Genet. 2018;49(5):457–60.

20. Yuan S, Norgard RJ, Stanger BZ. Cellular Plasticity in Cancer. Cancer Discov. 2019;9(7):837–51.

21. Chen C, Zimmermann M, Tinhofer I, Kaufmann AM, Albers AE. Epithelial-to-mesenchymal transition and cancer stem(-like) cells in head and neck squamous cell carcinoma. Cancer Lett. 2013;338(1):47–56.

22. Oshimori N. Cancer stem cells and their niche in the progression of squamous cell carcinoma. Cancer Sci. 2020;111(11):3985–92.

23. Baum B, Settleman J, Quinlan MP. Transitions between epithelial and mesenchymal states in development and disease. Semin Cell Dev Biol. 2008;19(3):294–308.

24. Dongre A, Weinberg RA. New insights into the mechanisms of epithelial-mesenchymal transition and implications for cancer. Nat Rev Mol Cell Biol. 2019;20(2):69–84.

25. Yang J, Weinberg RA. Epithelial-mesenchymal transition: at the crossroads of development and tumor metastasis. Dev Cell. 2008;14(6):818–29.

26. Babaei G, Aziz SG, Jaghi NZZ. EMT, cancer stem cells and autophagy; The three main axes of metastasis. Biomed Pharmacother. 2021;133:110909.

27. Dawood S, Austin L, Cristofanilli M. Cancer stem cells: implications for cancer therapy. Oncology (Williston Park). 2014;28(12):1101–7, 10.

28. Elkashty OA, Abu Elghanam G, Su X, Liu Y, Chauvin PJ, Tran SD. Cancer stem cells enrichment with surface markers CD271 and CD44 in human head and neck squamous cell carcinomas. Carcinogenesis. 2020;41(4):458–66.

29. Keysar SB, Le PN, Miller B, Jackson BC, Eagles JR, Nieto C, et al. Regulation of Head and Neck Squamous Cancer Stem Cells by PI3K and SOX2. J Natl Cancer Inst. 2017;109(1).

30. Ben-Porath I, Thomson MW, Carey VJ, Ge R, Bell GW, Regev A, et al. An embryonic stem cell-like gene expression signature in poorly differentiated aggressive human tumors. Nat Genet. 2008;40(5):499–507.

31. Maniotis AJ, Folberg R, Hess A, Seftor EA, Gardner LM, Pe’er J, et al. Vascular channel formation by human melanoma cells in vivo and in vitro: vasculogenic mimicry. Am J Pathol. 1999;155(3):739–52.

32. Lugassy C, Zadran S, Bentolila LA, Wadehra M, Prakash R, Carmichael ST, et al. Angiotropism, pericytic mimicry and extravascular migratory metastasis in melanoma: an alternative to intravascular cancer dissemination. Cancer Microenviron. 2014;7(3):139–52.

33. Massimini M, Romanucci M, De Maria R, Della Salda L. An Update on Molecular Pathways Regulating Vasculogenic Mimicry in Human Osteosarcoma and Their Role in Canine Oncology. Front Vet Sci. 2021;8:722432.

34. Thijssen VL, Paulis YW, Nowak-Sliwinska P, Deumelandt KL, Hosaka K, Soetekouw PM, et al. Targeting PDGF-mediated recruitment of pericytes blocks vascular mimicry and tumor growth. J Pathol. 2018;246(4):447–58.

35. Vartanian A, Stepanova E, Grigorieva I, Solomko E, Belkin V, Baryshnikov A, et al. Melanoma vasculogenic mimicry capillary-like structure formation depends on integrin and calcium signaling. Microcirculation. 2011;18(5):390–9.

36. Yang JP, Liao YD, Mai DM, Xie P, Qiang YY, Zheng LS, et al. Tumor vasculogenic mimicry predicts poor prognosis in cancer patients: a meta-analysis. Angiogenesis. 2016;19(2):191–200.

37. Liu ZL, Chen HH, Zheng LL, Sun LP, Shi L. Angiogenic signaling pathways and anti-angiogenic therapy for cancer. Signal Transduct Target Ther. 2023;8(1):198.

38. Guiraldelli GG, Prado MCM, de FLP, Leis-Filho AF, Kobayashi PE, Cury SS, et al. Pathways Involved in the Development of Vasculogenic Mimicry in Canine Mammary Carcinoma Cell Cultures. J Comp Pathol. 2022;192:50–60.

39. Nordio L, Fattori S, Vascellari M, Giudice C. Evidence of Vasculogenic Mimicry in a Palpebral Melanocytoma in a Dog. J Comp Pathol. 2018;162:43–6.

40. Strohmayer C, Klang A, Kummer S, Walter I, Jindra C, Weissenbacher-Lang C, et al. Tumor Cell Plasticity in Equine Papillomavirus-Positive Versus-Negative Squamous Cell Carcinoma of the Head and Neck. Pathogens. 2022;11(2).

41. Hanahan D, Weinberg RA. Hallmarks of cancer: the next generation. Cell. 2011;144(5):646–74.

42. Mani SA, Guo W, Liao MJ, Eaton EN, Ayyanan A, Zhou AY, et al. The epithelial-mesenchymal transition generates cells with properties of stem cells. Cell. 2008;133(4):704–15.

43. Armando F, Mecocci S, Orlandi V, Porcellato I, Cappelli K, Mechelli L, et al. Investigation of the Epithelial to Mesenchymal Transition (EMT) Process in Equine Papillomavirus-2 (EcPV-2)-Positive Penile Squamous Cell Carcinomas. Int J Mol Sci. 2021;22(19).

44. Miglinci L, Reicher P, Nell B, Koch M, Jindra C, Brandt S. Detection of Equine Papillomaviruses and Gamma-Herpesviruses in Equine Squamous Cell Carcinoma. Pathogens. 2023;12(2).

45. Rushton JO, Kolodziejek J, Tichy A, Nowotny N, Nell B. Clinical course of ophthalmic findings and potential influence factors of herpesvirus infections: 18 month follow-up of a closed herd of lipizzaners. PLoS One. 2013;8(11):e79888.

46. Hujanen R, Almahmoudi R, Karinen S, Nwaru BI, Salo T, Salem A. Vasculogenic Mimicry: A Promising Prognosticator in Head and Neck Squamous Cell Carcinoma and Esophageal Cancer? A Systematic Review and Meta-Analysis. Cells. 2020;9(2).

47. Hujanen R, Almahmoudi R, Salo T, Salem A. Comparative Analysis of Vascular Mimicry in Head and Neck Squamous Cell Carcinoma: In Vitro and In Vivo Approaches. Cancers (Basel). 2021;13(19).

48. Salem A, Salo T. Vasculogenic Mimicry in Head and Neck Squamous Cell Carcinoma-Time to Take Notice. Front Oral Health. 2021;2:666895.

49. Liu SY, Chang LC, Pan LF, Hung YJ, Lee CH, Shieh YS. Clinicopathologic significance of tumor cell-lined vessel and microenvironment in oral squamous cell carcinoma. Oral Oncol. 2008;44(3):277–85.

50. Valdivia A, Mingo G, Aldana V, Pinto MP, Ramirez M, Retamal C, et al. Fact or Fiction, It Is Time for a Verdict on Vasculogenic Mimicry? Front Oncol. 2019;9:680.

51. Chang YS, di Tomaso E, McDonald DM, Jones R, Jain RK, Munn LL. Mosaic blood vessels in tumors: frequency of cancer cells in contact with flowing blood. Proc Natl Acad Sci U S A. 2000;97(26):14608–13.

52. Folberg R, Hendrix MJ, Maniotis AJ. Vasculogenic mimicry and tumor angiogenesis. Am J Pathol. 2000;156(2):361–81.

53. Hendrix MJ, Seftor EA, Hess AR, Seftor RE. Vasculogenic mimicry and tumour-cell plasticity: lessons from melanoma. Nat Rev Cancer. 2003;3(6):411–21.

54. Hendrix MJ, Seftor EA, Seftor RE, Chao JT, Chien DS, Chu YW. Tumor cell vascular mimicry: Novel targeting opportunity in melanoma. Pharmacol Ther. 2016;159:83–92.

55. Seftor EA, Meltzer PS, Schatteman GC, Gruman LM, Hess AR, Kirschmann DA, et al. Expression of multiple molecular phenotypes by aggressive melanoma tumor cells: role in vasculogenic mimicry. Crit Rev Oncol Hematol. 2002;44(1):17–27.

56. Marelli-Berg FM, Clement M, Mauro C, Caligiuri G. An immunologist’s guide to CD31 function in T-cells. J Cell Sci. 2013;126(Pt 11):2343–52.

57. Saitoh M. Involvement of partial EMT in cancer progression. J Biochem. 2018;164(4):257–64.

58. Liao C, Wang Q, An J, Long Q, Wang H, Xiang M, et al. Partial EMT in Squamous Cell Carcinoma: A Snapshot. Int J Biol Sci. 2021;17(12):3036–47.

59. Franke WW, Schmid E, Osborn M, Weber K. Intermediate-sized filaments of human endothelial cells. J Cell Biol. 1979;81(3):570–80.

60. Miettinen M, Fetsch JF. Distribution of keratins in normal endothelial cells and a spectrum of vascular tumors: implications in tumor diagnosis. Hum Pathol. 2000;31(9):1062–7.

61. Kornfield TE, Newman EA. Regulation of blood flow in the retinal trilaminar vascular network. J Neurosci. 2014;34(34):11504–13.

62. Stratman AN, Malotte KM, Mahan RD, Davis MJ, Davis GE. Pericyte recruitment during vasculogenic tube assembly stimulates endothelial basement membrane matrix formation. Blood. 2009;114(24):5091–101.

